# Balanced contractility and adhesion drive polarization in a minimal elastic actomyosin network

**DOI:** 10.1101/2025.08.26.672435

**Authors:** Zeno Messi, Franck Raynaud, Nathan Goehring, Alexander B. Verkhovsky

## Abstract

Polarization of migrating cells involves chemical and mechanical interactions of signaling networks, cytoskeleton, plasma membrane, and substrate adhesions. Still, it is not fully understood which mechanisms and components are sufficient for symmetry breaking, and if they work independently or together. Here, we use a discrete active network model to investigate if and how an elastic cytoskeletal network is capable of breaking symmetry solely through mechanical interactions. Our minimal model consists of elastic bonds, attractive force dipoles, and force-sensitive anchor points, initially distributed uniformly and subject to simple turnover rules. We find that these features are sufficient to produce different cell behaviors, and, remarkably, to drive symmetry breaking and directed (polarized) motion. Network behavior was primarily determined by the turnover rate of anchor points, which, itself, is a function of the ratio between dipole force and the threshold force required for anchor removal. Directional motion emerged at intermediate turnover rates, at which tension in the network accumulated through several turnover cycles before eventually exceeding the adhesion removal threshold locally at the edge, mirroring our recent experimental findings on correlation of the traction force with protrusion-retraction transitions in the cell [1, 2]. At high turnover rates, forces were unable to build up to sufficiently high levels, while at low turnover rates, anchors hinder motion. These results demonstrate how directed motion can emerge as an intrinsic property of a simple mechanical network, independently of external cues or complex signaling networks. Given the concordance between this model and recent experimental findings, we suggest that polarization by contraction-adhesion dynamics could be a fundamental emergent behavior of actin-myosin networks.

**Author summary:** Cells often need to move, for example, during development, wound healing, or immune responses. To do so, they must first decide where their “front” and “back” are, a process known as polarization. Most explanations for this behavior focus on complex chemical signaling inside the cell. In our work, we asked a simpler question: could mechanical forces within a cell be sufficient to make it polarize and move without an external cue? To explore this idea, we developed a computational model of a simplified cell made only of elastic connections, contractile forces, and attachment points to its surroundings. We started with a completely uniform system, without any built-in direction or external guidance. Surprisingly, we found that this minimal mechanical setup could spontaneously develop a front and a back and begin moving persistently. We proposed that the key mechanical factor controlling this behavior was the rate at which the attachment points break under mechanical load. When this process occurred at an intermediate rate, forces built up unevenly, leading to detachment and forward motion, similar to what is observed *in-vitro*. Our findings suggest that cell polarization and movement can emerge spontaneously from mechanical properties alone, highlighting an important and overlooked role for mechanics in cell behavior.

## Introduction

Migrating cells can break symmetry, forming a structurally and functionally distinct protrusive front and retractive rear, and maintain this polarization during persistent motion. Polarization is a complex process involving the interaction of multiple components, including signalling networks, the plasma membrane, the cytoskeleton, and adhesions [3–9]. Although decades of biochemical and mechanical studies [10, 11] have illuminated the properties and dynamics of individual players such as actin filaments, myosin motors, adhesion complexes, and the membrane, the integrative physical mechanisms initiating and subsequently stabilizing cell polarity remain unresolved. Polarization can occur in response to various external signals, such as chemical gradients, mechanical stimuli, and electric fields [12–17]. However, equally striking is that cells can polarize spontaneously, without an imposed directional cue, implying that symmetry breaking is an intrinsic property of the motile apparatus itself [18, 19]. It could be initiated by localized protrusion at the prospective cell front, but also by the retraction at the prospective rear [20, 21]. This suggests that polarization may occur via multiple pathways and that the mechanisms responsible for breaking symmetry may be distinct from those responsible for sensing external cues. Mathematical and computational models are pivotal to decipher this complexity by reformulating the problem in terms of simpler, individual, and testable processes that can reveal whether—and under what conditions—those components would spontaneously organize themselves to create front–rear polarity [22, 23]. Several models attempted to include multiple, if not all, players of polarization processes, describing polarization in terms of feedback loops between the signaling networks, membrane, and the cytoskeleton [24–26]. These include reaction-diffusion dynamics of cytoskeletal regulators such as small GTPases [27], feedback between membrane tension or curvature and actin dynamics [28, 29], coupling between polarity and motion through the flow of the actin network [30], as well as combinations of these mechanisms [26, 31].

Models with a narrower focus relied on purely cytoskeletal mechanics. Discrete elastic model of actin-myosin network investigated the effect on nonlinear actin elasticity on force transmission in a static isotropic configuration [32]. To investigate polarization, other studies utilized description of actin-myosin network as viscoelastic continuum [33–35]. Several types of feedback relationships and their combinations were considered: non-linear feedback between actin flow and adhesions, force feedback from the outer boundary of the system, and competition between contractile and protrusive actin populations for a limited monomer pool. However, these continuum models were not well suited to analyse initial stages of polarization: they required manual introduction of either a significant initial asymmetry or very large fluctuations (correlation length and decay time comparable to dimensions of the whole simulation [33]

Here, we take a different approach: we extend discrete elastic model of actin-myosin network to dynamical case by adding simple rules of turnover. Most importantly, we introduce an intuitive rule for adhesion turnover: when adhesions experience a force above threshold, they break. This is inspired by our previous studies that demonstrate that increased traction forces at the cell periphery coincide with the cell edge switching from protrusion to retraction, eventually resulting in polarization [1, 2]. In our minimalistic system of elastic bonds, force dipoles, and fixed substrate anchors any feedback relationships and eventual symmetry breaking represent true emergent properties of local mechanics. We demonstrate that, provided the optimal balance between contractile and adhesive elements, this system can self-organize and break symmetry through solely mechanical interactions, without any feedback from chemistry, outer boundary, or mass conservation constraints.

## Material and methods

### Cell computational model

#### Description and initialization

Our computational model is composed of three elements: elastic bonds, attractive dipoles, and anchors, which represent, respectively, actin filaments, non-muscle myosin II mini-filaments, and cell-substrate adhesion. Elastic bonds and dipoles connect the vertices of an underlying regular hexagonal lattice. They are deformable elements that can be stretched, compressed, or bent at their midpoint to create a hinge. Bonds have a rest length 2*ℓ*_0_. Attractive dipoles can form between any pair of vertices within a distance of a bond. It is not required that a bond underlies a dipole. Anchors are fixed vertices attached to the substrate.

The system is initialized with a uniform random distribution of these three components inside a disk with a radius of 20 bonds. Bonds, dipoles and anchors have respective densities *ρ, ρ*_*d*_, and *ρ*_*a*_. After the system is initialized, it is allowed to find an energy minimum.

#### Energy and model units

To compute the energy in the system, we model actin filaments using the Worm-Like-Chain (WLC) model like in [2] and [32]. In the WLC [36], filaments are considered a chain of inextensible segments which relative orientation can vary. The filaments are characterized by their persistence length *ℓ*_*p*_ which can be regarded as the filament length at which significant bending fluctuations occur. In this framework, the bending energy a filament of length 2*ℓ*_0_ with curvature *R* reads

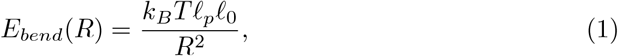

where *k*_*B*_*T* is the thermal energy. Assuming bending is uniform, we can substitute the curvature radius and rewrite the bending energy as a function of the angle *ϕ* between the two ends of the segment as

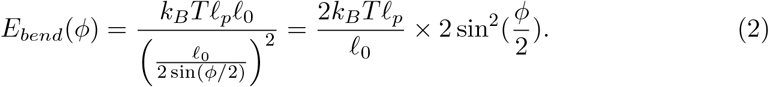

In the model, bonds can bend in their middle to form a hinge. On top of that, consecutive bonds that are initially aligned also form hinges and contribute to the bending energy of the system. Connected bonds that are not initially aligned can change relative orientation freely. The energy required to fully bend a hinge is

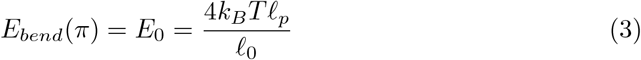

and the typical force for bending a filament is

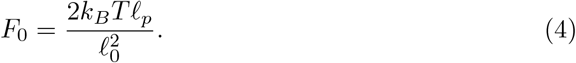

In the WLC model, individual segments are inextensible. However, thermal fluctuations result in a contraction of the filament end-to-end distance and the filament can be stretched if tension is applied. Following [36], in the linear regime, the stretching energy of a segment of length *ℓ* and rest length *ℓ*_0_ is

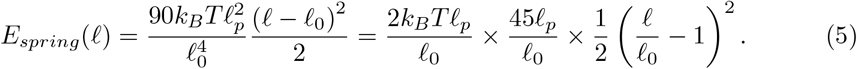

The two other contributions to the energy of the system are the dipole and substrate deformation. The attractive force between dipoles is constant and thus the energy associated with dipoles is

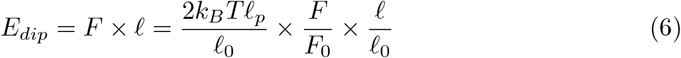

where *F* is the dipole force and *ℓ* is the distance between the nodes forming the dipole. Finally, the energy of the deformation of the substrate depends on the distance *d* between the current position of the anchor and its initial position and thus reads

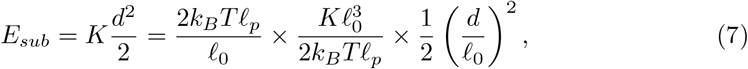

where K is an elastic constant and d is the distance between the current position of the anchor and its initial position. We can write the full Hamiltonian:

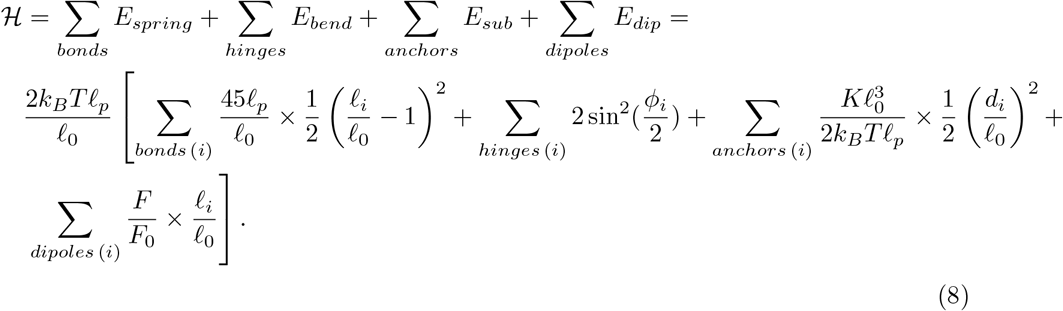

To work with dimensionless parameters, we set the factor of the last equation

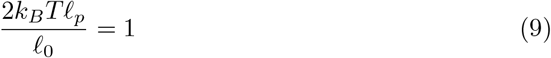

and we define the dimensionless parameters:

- bond spring constant

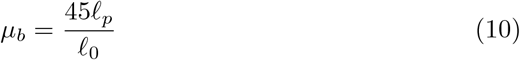
- dipole force magnitude

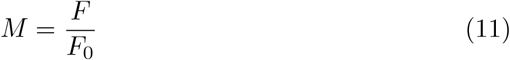
- substrate spring constant

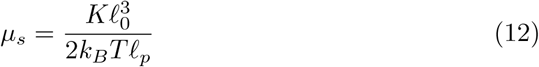

Finally, we set the rest length of the segments

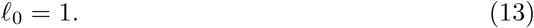

With these definitions, the full Hamiltonian becomes

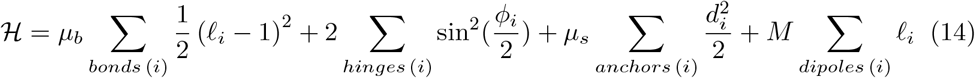

We can now identify biologically relevant ranges for our model. The mesh size can be identified to the length of myosin bipolar minifilaments which have a typical length of 0.1 − 1*µm* [37]. The rest length of actin filaments is *ℓ*_*p*_ ~ 10*µm* [38, 39](other refs?).

This gives a range for the bond spring constant

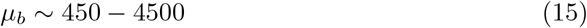

In this work we take an intermediate value for this parameters and set *µ*_*b*_ = 1024. To get the force required to bend a hinge we plug the numbers in equation 4

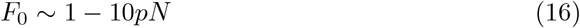

which is consistent with experimental measurements [40, 41]. In our simulations, when a single dipole is not capable to bend a bond, the system simply grows. Therefore, we limited our analysis to conditions where single dipoles can bend bonds. We set the range of dipole force magnitudes between 1 and 16. Finally, for the elastic modulus of the substrate, we set its value to be very high, we set

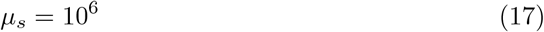

#### Implementation of the dipole force and energy in the model

For numerical stability, we chose to set a continuous cut-off on the force at short distances in the implementation of the model. The force acting on a vertex of a dipole reads

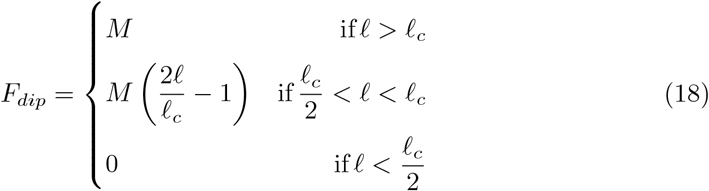

where *ℓ* is the dipole length, *M* is the dipole force magnitude parameter, and *ℓ*_*c*_ is the cut-off length. The energy associated with the force dipole is

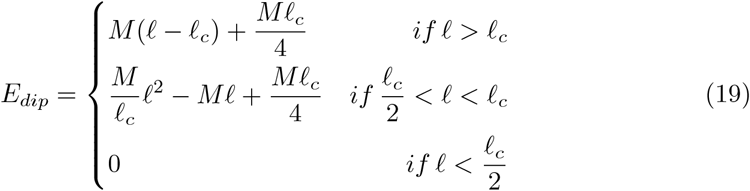

The parameters of the model are listed in Table 1. They can be classified into several groups: structural, including the bond, anchor, and dipole densities; energetic, encompassing the dipole force and bond’s spring constant; and parameters related to the turnover and pruning of the network, such as the anchor removal threshold and the protrusion rate. Values of some of the structural and energetic parameters were selected based on the knowledge about properties of the actin-myosin network and previous modeling work [2, 32]. Here, we focused on the variation of parameters defining the turnover of anchors: the anchor density, the anchor removal threshold and the dipole force magnitude. We found that these parameters were critical for the evolution of the system and gave rise to qualitatively distinct phenotypes. Changing other parameters only shifted the boundaries in the parameter space but did not produce additional phenotypes (not shown).

**Table 1.** Model parameters.

| Name | Description | Range |
| --- | --- | --- |
| $\rho$ | lattice density | 0.7 |
| $\rho_a$ | anchor density | 0.05 – 4% |
| $\rho_d$ | dipole density | 0.3 |
|  | max bonds per pair of nodes | 10 |
|  | max dipoles per pair of nodes | 10 |
| $\ell_0$ | bond rest length | 1 |
|  | energy required to fully bend a bond | 2 |
| $\mu_b$ | bond spring constant | 1024 |
| $\mu_s$ | substrate spring constant | $10^6$ |
| $T_a$ | anchor removal threshold | 0 – 16000 |
| $M$ | dipole force magnitude | 0 – 16 |
| $P$ | protrusion rate | 4 |
| $d_{merge}$ | node merging distance | 0.02 |
| $\alpha_{rem}$ | hinge removal angle | $\cos^{-1}(0.8) \approx 37^\circ$ |
| $\ell_c$ | dipole force cut-off length | 0.01 |
| $D$ | removal threshold gradient strength | 0 – 4 |

#### Pruning

Compared to previous studies [2, 32, 36], our computational cell model evolves through hundreds of consecutive energy minimization procedures. However, as no steric effects are considered in the model, it is necessary to remove some components to avoid spurious aggregation of material that would perturb further computation and minimization of the Hamiltonian.

1. Using equation (7), the force applied on each anchor is computed. The anchors withstanding a force higher than a given threshold *T*_*a*_ are removed;
2. The hinges that are bent below an angle lower than *α*_*rem*_ are removed;
3. The vertices that are closer than a threshold distance are merged into a single node:
  - Nodes are iteratively grouped by pair and are merged when the distance between the nodes is smaller than a threshold distance *d*_*merge*_.
  - Merged nodes are replaced by a new node located at their barycenter carrying all the bonds and dipoles of the pair. If one of the two merged nodes was an anchor, the new node is also an anchor.
  - We fixed a maximum of 10 bonds attached to a node and 10 dipoles per pair. After merging, whether there were more than 10 dipoles attached to a pair of nodes, the exceeding dipoles are reassigned at random to other pairs of nodes. The exceeding bonds are removed from the system
4. Nodes that become disconnected, that do not belong to the largest connected component, are also removed.

#### Outline measurement and network growth

The final step consists in adding new network at the edge of the system. First, the system boundary is found with a method derived from [42]. In brief, we consider all nodes that remain after the pruning procedure and compute the Delaunay triangulation of this set of nodes (if needed, fictive points are added at a distance of 0.01 around nodes to avoid singular edges in the triangulation). The boundary edges of the convex hull are directly identified from the triangulation, as they correspond to edges belonging to a single triangle; however, it is necessary to account for concavities. Edges of the convex hull are iteratively removed, and the convex hull recomputed after each removal, until no edges of the initial set with a length greater than 5 are present. This method [42] enables outlines with concavities and ensures that there are no holes within the shape. Then, each point of the outline is translated normally by a distance *P* to mimic actin protrusion at the edge of the cell. New material, comprising bonds, dipoles, and anchors, is added between the shifted and original outlines. The density of the new network components is the same as at initialization, new network bonds are then connected to the existing network if they are at a distance from the original network shorter than the rest bond length. The simulation step is done, and the minimization can be carried out once again.

#### Shuffling of anchors and dipoles

To test the effect of loss of anchor and/or dipole polarity, we considered a restarted the simulation from a well polarized initial state randomizing anchors and/or dipoles. We simulated the system for 50 time steps and compared the trajectories and outlines obtained in the different conditions (control, shuffle dipoles, shuffle anchors, shuffle both).

#### Simulations with external cue

The external cue is simulated by varying the anchor threshold in the system. For any anchor, the effective removal threshold *T*_*eff*_ is given by a baseline *T*_*a*_ shifted by a constant multiplied by the relative distance between the x-coordinate of the anchor and the center of the system

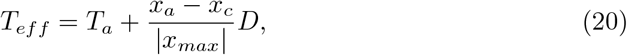

where *D* is the gradient strength, *x*_*a*_ and *x*_*c*_ are the positions of the anchor and the center respectively. The distance is normalized by the distance to the anchor furthest from the center of the system.

#### Model implementation and data analysis

The model was implemented in a C++ program and the data was analyzed with C++ and python programs. For energy minimization, the GNU scientific library implementation of the BFGS algorithm was used [43]. Computational geometry for finding and expanding the boundary of the system was carried out with the CGAL library [44].

### Quantification and statistical analysis

#### Kymographs

To obtain linear kymographs, we generated raster images of the system network, anchors, and dipoles at each simulation time step. The resulting image sequences were imported into Fiji. For each sequence, we manually drew a line through the approximate center of the system, and used Fiji’s Reslice tool [45] to produce the kymograph. In polarized systems, the kymograph axis was aligned approximately with the front–back axis. approximately to the front-back axis.

To construct circular kymographs, anchors and dipoles were radially projected onto a circle centered on the system centroid. At each simulation step, the system was divided into 30 circular sectors radiating from this centroid (defined as the centroid of the system outline). The number of anchors (resp. dipoles) in each sector was counted and normalized to the maximum value at that time step. The normalized values were then color-coded to generate a single line of the kymograph.

#### Network flow

Network flowwas quantified as the average displacement of nodes before and after the minimization procedure.. The system was divided into a 40 × 40 grid, scaled to the system size (with the number of squares fixed and their dimensions varying accordingly). The vectorial displacements of all lattice nodes were averaged in each square and represented by an arrow. Arrow length and color encode the magnitude of the average displacement.

#### Anchor lifetime

To quantify anchor lifetime, we counted how many how many time steps anchors existed before being removed. Every anchor in the system was followed from creation to removal or to the end of the simulation. If an anchor was not removed at the end of the simulation, its current lifetime was taken into account.

#### Trajectory gyration radius

To quantify the persistence of motion, we measured the gyration radius of trajectories. Each trajectory was treated as a point cloud, and its gyration radius was computed as the root-mean-square distance of all trajectory points from the cloud’s center of mass

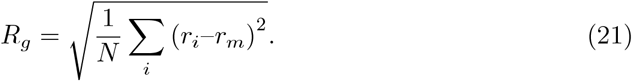

Compared to persistence measures based on end-to-end distance, the gyration radius incorporates the entire trajectory, and remains meaningful even when trajectories include pauses, loops, or changes of direction.

#### Force maps

Force maps were generated from raster images of the forces acting on anchors. The system divided into a grid of 517×517 of pixels. For every pixel in the grid, the assigned force was the average of contributions from all nearby anchors, up to a distance of 40 length units. This approach does not represent a physical force field, but rather provides a qualitative visualization of the spatial distribution and relative magnitude of forces. In addition to the force image, we created three other images for trajectory, system outline, colormap. All four images were imported into Fiji, where the force image was blurred and then blended with the other layers to produce the final visualization.

#### Radial histograms

For simulations with an external cue and migrating phenotype, radial histograms were generated from the angular distribution of the end-to-end vector, pooled across 20 simulations per parameter set. For the radial phenotype, histograms were instead computed from the angular distribution of all displacements recorded during the simulations. In both cases, bins represent a 30 degrees angle.

#### Persistency score

To compare persistency between cells of different size, we define a persistency score *S*_*pers*_ as follows: first, we calculate the mean squared displacement MSD(*τ*) for different lag values *τ* :

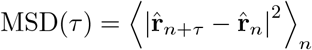

Here 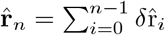 and 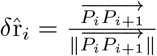 is a normalized vector pointing along the direction of two successive positions of the geometrical cell center *P*_*i*_ and *P*_*i*+1_. In this representation, the MSD of ballistic motion (maximal persistence) and that of two-dimensional random motion (no orientational persistence) bound the MSD of any simulated trajectory. For a ballistic motion, we have MSD_ballistic_(*τ*) = *τ*^2^, while for a random motion MSD_random_(*τ*) = *τ*. The score *S*_*pers*_ is calculated as:

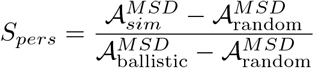

where 𝒜^MSD^ represents the area under the MSD curves for the simulated cells, ballistic motion, or random motion. Consequently, if the cell trajectory is a straight line, *S*_*pers*_ = 1, whereas if the trajectory is random, *S*_*pers*_ = 0.

## Results

### A mechanical model of self-organized cell migration

The core of the model is based on a network of elastic elements, bonds, connected via nodes, effectively representing a minimal actin network [2, 32]. Similar to actin filaments, bonds can undergo stretching, compression, and bending when subjected to force. To mimic asymmetric elasticity of actin filaments, parameters of the model are set such that bending requires significantly less force than stretching or compression (see Material and Methods). The two additional mechanical elements are attractive force dipoles that exert contractile force between neighboring pairs of nodes, reflecting myosin, and anchors, which take the form of fixed-position nodes and mimic substrate attachment sites. Importantly, the anchors are force sensitive – if the applied force exceeds a threshold, the anchors are removed.

To explore the behavior of this system, networks were initialized on a hexagonal lattice with random distributions of bonds, dipoles, and anchors (Figure 1A). Networks were then evolved as follows. They were allowed to deform to find a conformation of minimal energy, and anchors experiencing force above threshold were removed. The network was then pruned. Briefly, highly bent bonds were removed, and pairs of nodes closer than a threshold distance were merged. Merging could result in multiple bonds and dipoles connecting the same nodes. If there were more than 10 dipoles and/or bonds connecting a pair of nodes, excess dipoles were reassigned randomly, and excess bonds were removed (Figure 1B, see Methods for details). Finally, the edge of the system was extended isotropically, mimicking actin polymerization, with new bonds, dipoles and anchors, and the network was allowed to deform again to find a new conformation of minimal energy (Figure 1C). This sequence was repeated at each step of the simulation. The simulations were stopped after 400 steps or if the system exceeded a threshold area of 100000 model units (see Methods for details).

**Fig 1.**
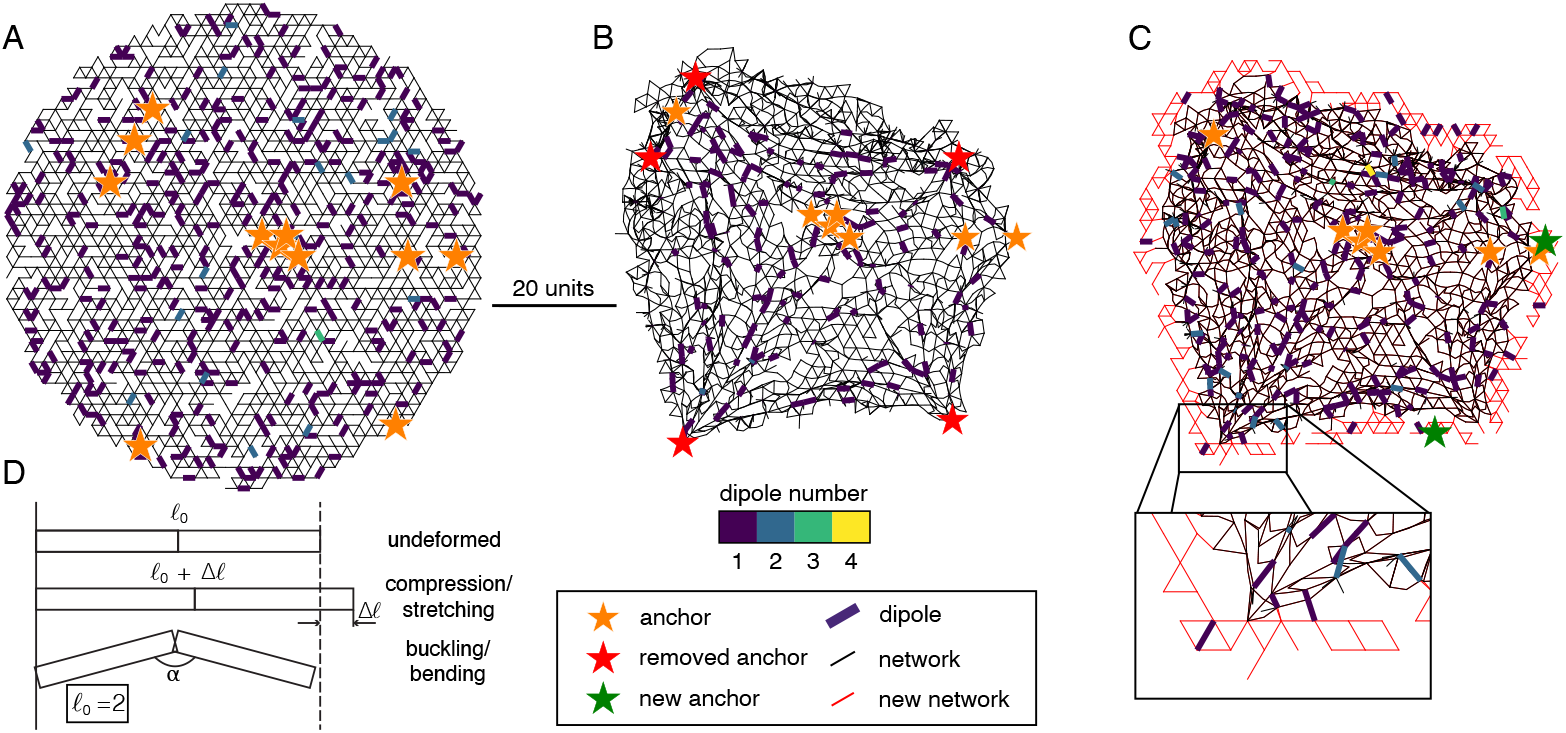
Model description and dynamical rules. A-C) State of the system at different stages of a simulation step. Bonds and anchors are represented as black segments and orange stars, respectively. The number of dipoles connecting the same network node are color coded from dark blue (low number) to yellow (high number). Scale bar represents 20 model distance units. A) State of the system at initialization, in a circle of 40 units radius with uniform distribution of bonds, anchors and dipoles. B) State of the system after energy minimization. After minimization, anchors that are subjected to a force exceeding force threshold (in red) are removed. Only dipoles with a non-zero extension are displayed. C) State of the system after merging of neighboring nodes, pruning of highly bent hinges, and addition of new bonds (red) and anchors (green). D) Description of the three modes of bond deformation: stretching, compression and buckling under load.

### Assessing system size and motion across model parameters

As previously described, the system evolves through an iterative process: isotropic addition of new material along the perimeter, followed by energy minimization. We first investigated the mean area and its temporal evolution across an ensemble of over 150 distinct parameter sets (Figure 2A). Since material addition in our model is unconstrained by design, we anticipated unbounded area growth. We observed this behavior in half of the parameter sets (Figure 2A insets). While unconstrained growth is of limited biological relevance, it is notable that the remaining parameter sets produced areas that stayed below a prescribed threshold. Most cells, however, maintained their area below this limit and stabilized at a stationary area over time, rather than merely growing more slowly (Figure 2A insets).

**Fig 2.**
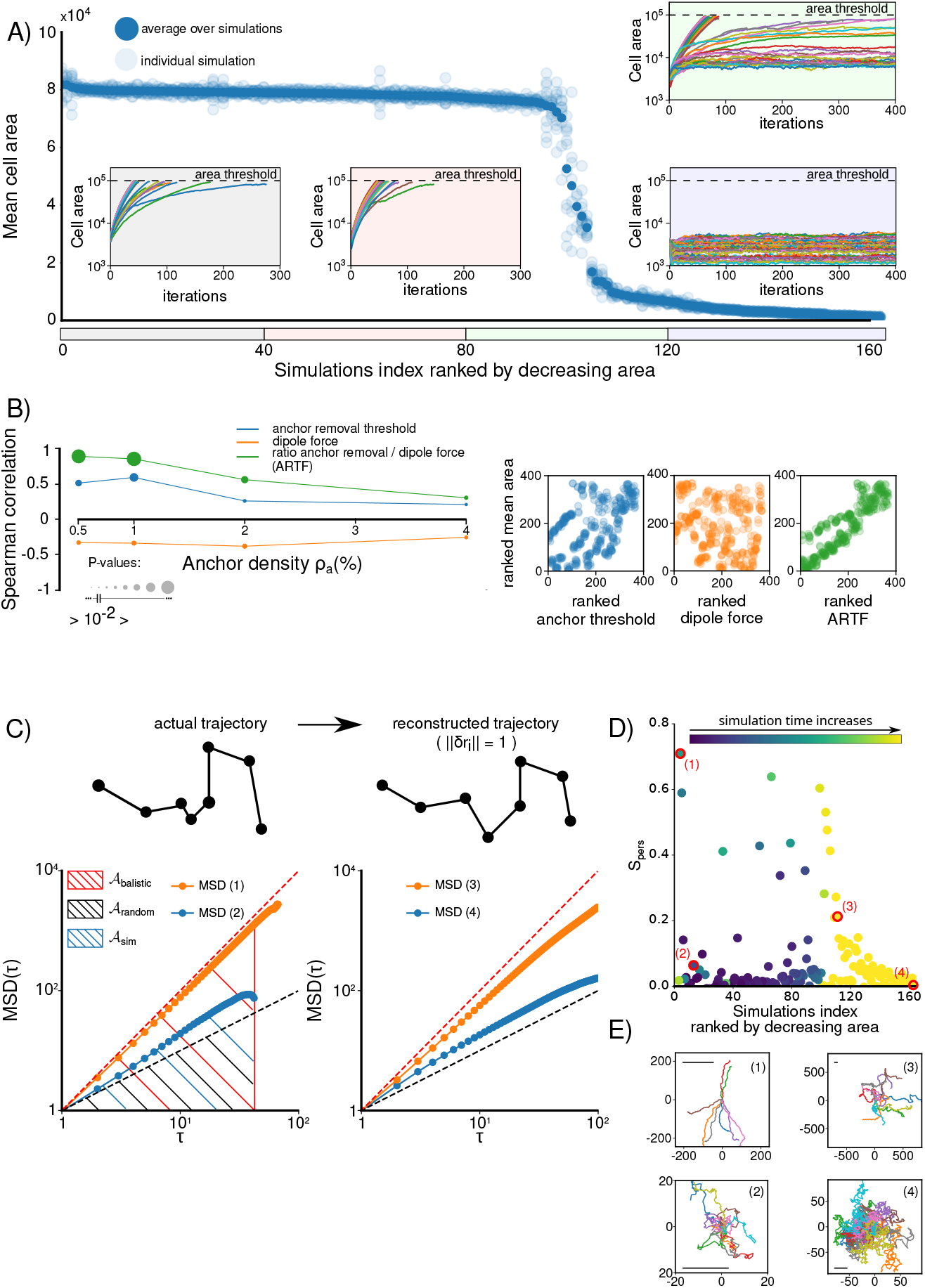
Model behavior. A) Mean cell areas plotted against the simulation index, ranked by decreasing area. Transparent blue points represent individual simulations, while solid blue points denote the average over identical parameter sets. For each simulation, the mean area is computed over the final third of the simulation duration. Insets show the temporal evolution of cell area, averaged across identical parameter sets; each inset corresponds to a group of 40 distinct parameter sets. B) (Left) Spearman correlation coefficients between cell area and anchor removal threshold (blue), dipole force (orange), and ARTF (green) as a function of anchor density. (Right) Scatter plot of ranked mean cell area versus ranked anchor removal threshold (blue), ranked dipole force (orange), and ranked ARTF (green) for a fixed anchor density *ρ*_*a*_ = 0.5%. C) Mean Squared Displacement (MSD) of normalized trajectories. The upper panel sketches the construction of a normalized trajectory. The lower panels display MSD(*τ*) for representative large cells (left) and small cells (right). The bounds for ballistic motion (MSD_*ballistic*_, red) and random motion (MSD_*random*_, black) are shown for reference. D) Scatter plot of the persistence score *S*_*pers*_, ordered by decreasing mean cell area. The color scale indicates simulation duration, from short (dark violet) to long (yellow). Four red circles are representative cases: large cells with high (1) and low (2) *S*_*pers*_, and small cells with high (3) and low (4) *S*_*pers*_. These specific cases correspond to the MSD curves shown in panel C. E) Cell trajectories corresponding to the four cases marked in D (1–4). Trajectories are non-normalized. Each trajectory represents one realization of a given set of parameters. The scale bar represents one typical cell radius for (1), (2), and (4); for (3), the scale bar represents 1*/*8 of the cell radius. The typical cell radius is calculated as 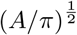, where *A* is the mean area.

We then investigated which parameters and parameter combinations determine cell size. For each simulation, we first assessed the associations between cell area and both the anchor threshold and the force dipole using Spearman rank correlations (Figure 2B). Across all anchor density values, cell area showed a significant positive correlation with anchor threshold and a significant negative correlation with force dipole. Accordingly, the ratio of anchor removal threshold to dipole force (ARTF) exhibited the strongest association with cell area, an association that weakened as anchor density increased.

Next, we examined the temporal characteristics of cell center displacement. Monitoring displacement alone is insufficient for comparing cells of different sizes, as two simulated cells can exhibit vastly different net displacements solely due to size differences. A more informative approach is to assess the persistence of the motion. To enable comparison across cells of varying sizes, we defined a persistence score *S*_*pers*_ based on the mean squared displacement (MSD) of a normalized trajectory of the cell center (see Material and Methods). This metric facilitates the comparison of persistence across all parameter sets independent of cell size and instantaneous velocity, avoids explicit curve fitting for MSD(*τ*), and penalizes persistence at short time scales.

Figure 2C displays individual MSD curves for representative cells with large and small areas, while Figure 2D plots the *S*_*pers*_ values for all simulations, ordered by decreasing cell area. Generally, large cells exhibit low persistence scores; *S*_*pers*_ increases as cell area decreases, but eventually trends toward 0 for the smallest cells. Inspection of individual trajectories (Figure 2E) reveals distinct phenotypic behaviors: small cells exhibiting either erratic or persistent motion, expanding but stationary cells, and larger cells that appear more persistent. In this latter regime, cells simultaneously grow and undergo translocation of their geometric centers; however, the actual displacement remains small relative to the typical cell size. The following section provides an in-depth characterization of these distinct behaviors.

### Types of system behaviour dependent on anchor turnover parameters

The ARTF ratio and the anchor density determined how often the anchors were removed by the forces built in the network. Phenotypes resulting from variation of these parameters could be classified into 5 types.

#### Erratic and centripetal

For low ARTF ratios (0 to 3) nearly all anchors are removed at each step. Resulting configurations of the network, evolution of system outlines, and motion and distribution of the elements are illustrated in Figure 3. If the anchor density parameter is low, the system shrinks from its initial size and remains small. At any simulation step, there are only a few anchors that are removed immediately, leading to erratic contractions and jittering motion of the system (Supplementary Movie S1). Higher anchor density leads to a larger system with anchors deposited and subsequently removed in a band at the periphery, resulting in a more organized contraction in a centripetal direction and radial organization of the network (see network organization and vector maps of flow (Figures 3G and 3L) and Supplementary Movie S2). Both erratic and centripetal systems are characterized by concentration of dipoles near the system center and absence of anchors anywhere except the very periphery (Figure 3). To further illustrate the dynamics of these systems, we generate kymographs of two types: linear and circular. Linear kymographs along an arbitrary line (Figures 3E and 3K) show contractions at the periphery as well as general erratic motion of the system and demonstrate that anchors at the periphery of the system are short-lived. To create circular kymographs, we radially projected anchors and dipoles at each step onto a circle drawn around the system and displayed linearized circles side by side in a time sequence (Figures 3C and 3I). Circular kymographs of both erratic and centripetal systems show that anchors and dipoles are distributed in nearly isotropic fashion over the time sequence. The centripetal system is reminiscent of cells with weak adhesions displaying strong isotropic retrograde flow [46, 47].

**Fig 3.**
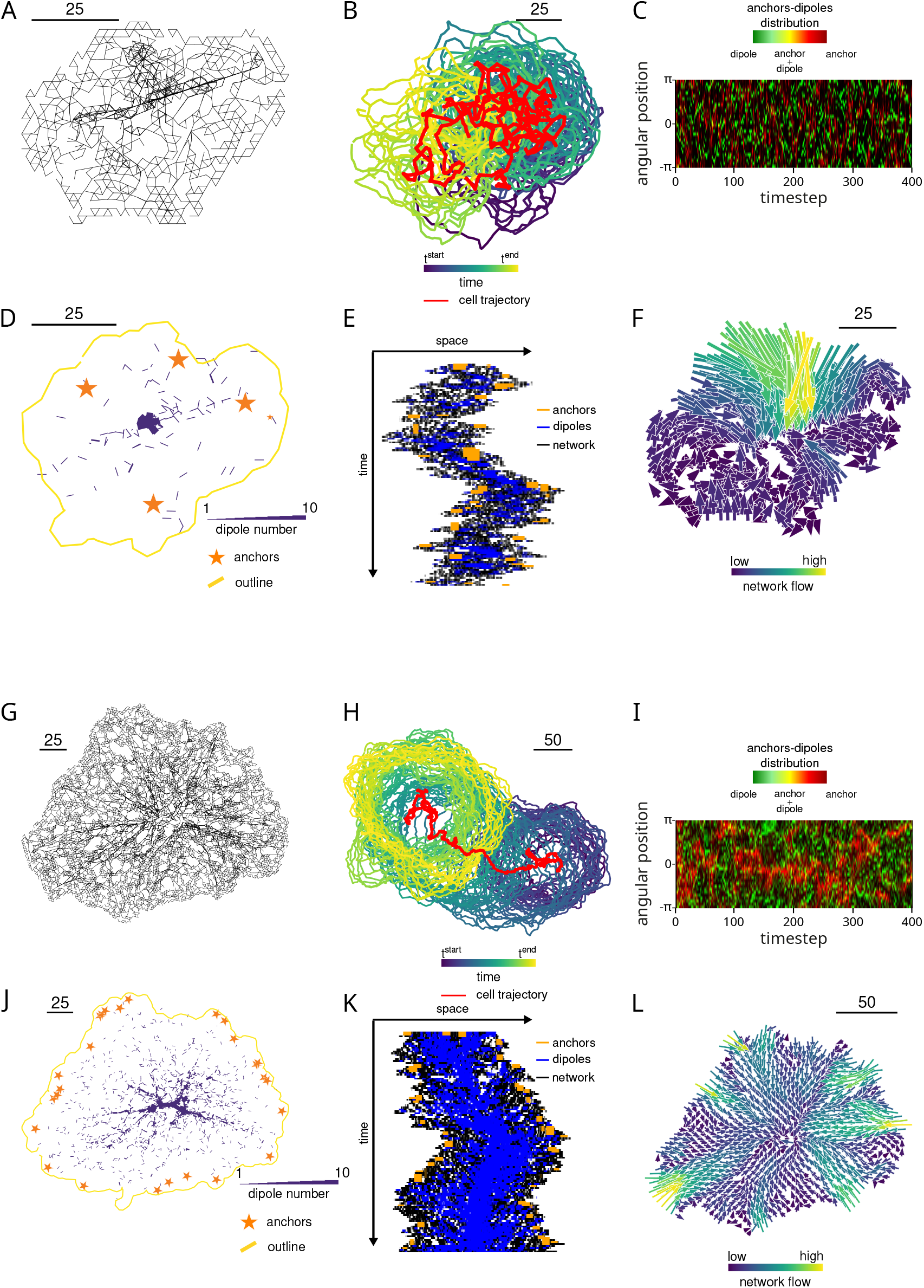
Low ARTF ratio leads to erratic motion and fast centripetal flow. Features and characteristic movement of systems with low ARTF ratio, and low anchor density (A-F) or high anchor density (G-L). A,G) Network configuration. B,H) Time evolution of the cell contour over 400 iterations. The position of the centroid is drawn in red. The system outline is shown every 10 steps, color coded for time from dark blue (first iteration) to yellow (last iteration). C,I) Circular kymograph of dipoles (green) and anchors (red) (see methods). D,J) Spatial distribution of anchors (orange stars) and dipoles (blue) with the system outline (yellow). The width of dipoles represents the number of overlapping dipoles. E,K) Kymograph along a line going through the center of the system. Bonds are represented in black, dipoles in blue, and anchors in orange. F,L) Network displacement before and after minimization. Amplitude of the displacement is color coded from dark blue (low flow) to yellow (high flow). The scale bar is given in model distance units.

#### Isotropic and anisotropic growth

For very high ARTF ratios (10 and above), the force generated in the network never reaches the anchor threshold, and anchors are not removed irrespective of their density. This phenotype is characterized by round systems undergoing isotropic growth. Growth is unlimited because our model does not feature mass conservation. These systems never move; their center of mass remains within the area occupied by the first iteration (Supplementary Movie S3, Figures 4A-F). Only the peripheral region of the system, where new network was recently added, undergoes some rearrangement and pruning (Figure 4F, Supplementary Movie S3). The rest of the system is frozen (Figures 4E and 4F). Rearrangements at the periphery during growth of the system later become frozen in the form of small alternating regions of high and low density of bonds and dipoles with anchors spread uniformly over the area. Overall distribution of all elements in the system is uniform and isotropic (Figure 4C). Such systems with narrow peripheral dynamical region and frozen bulk are similar to cells spreading in the conditions of impaired contractility.

**Fig 4.**
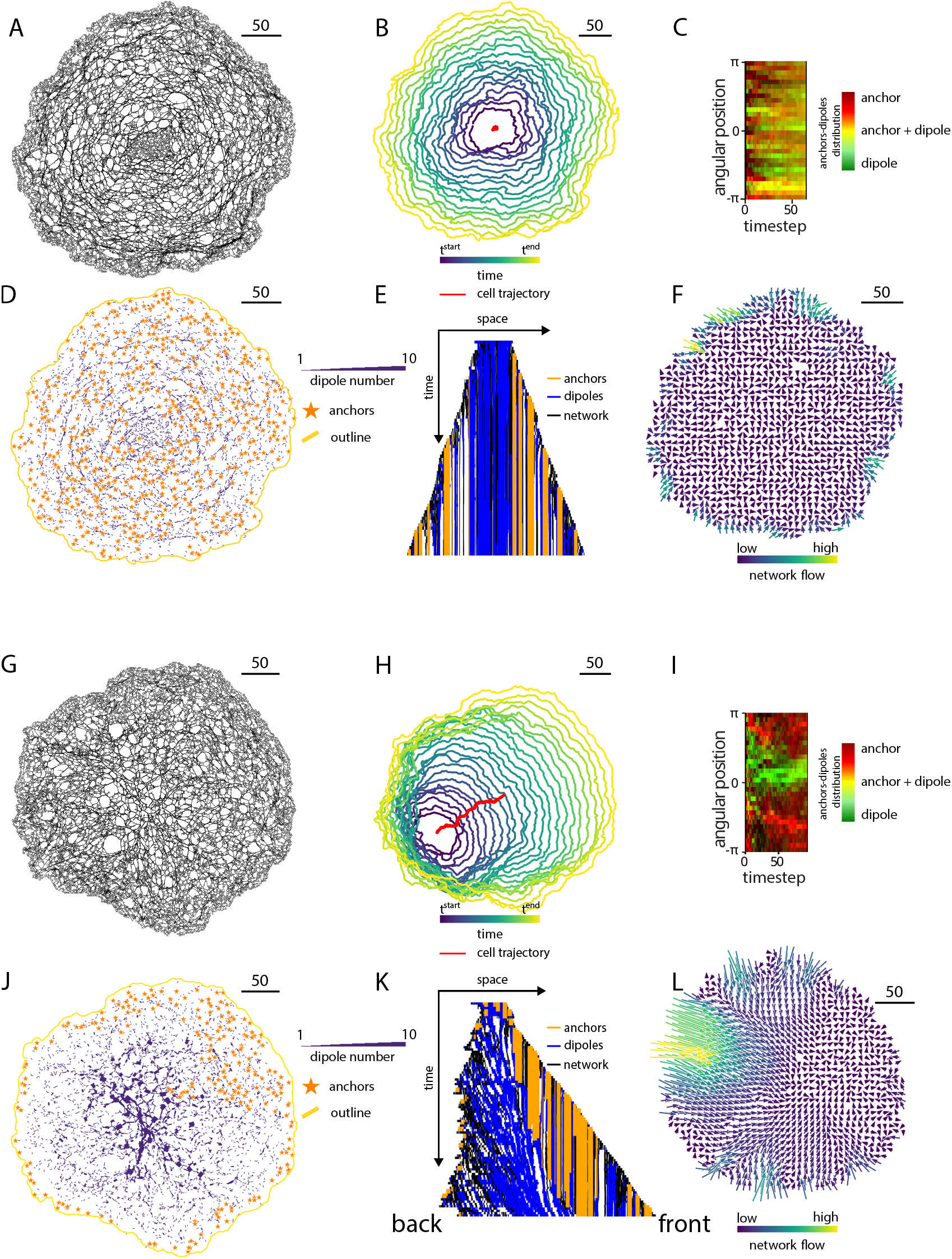
High ARTF ratio leads system growth. Features and characteristic movement of systems with high ARTF ratio. A,G) Network configuration. B,H) Movement of the system throughout the simulation (400 steps). The position of the centroid is drawn in red. The system outline is shown every 10 steps, color coded for time from dark blue to yellow (viridis palette). C,I) Circular kymograph of dipoles (green) and anchors (red) (see methods). D,J) Distribution of anchors (orange stars) and dipoles (blue) with the system outline (yellow). The width of dipoles represents the number of overlapping dipoles. E,K) Kymograph along a line going through the center of the system. Network is represented in black, dipoles in blue, and anchors in orange. F,L) Network displacement before and after minimization. Amplitude of the displacement is color coded from dark blue (low flow) to yellow (high flow). The scale bar is given in model distance units.

Lowering the ratio between anchor removal threshold to dipole force (typically, between 5 and 10) results in systems that also display unlimited growth, but the growth now has a preferred axis (Figure 4, Supplementary Movie S4). These systems do not move in the sense that the resulting trajectory never leaves the area occupied by their original contour, but they display a striking polarization. One side of the system grows steadily, while the other remains largely in place (Figure 4H). At the growing side, the network is frozen in the bulk and the anchors remain throughout the simulation, while the opposite side undergoes cycles of anchor removal and contraction and displays a flow of the network away from the periphery (Figures4K and 4L). Thus, the edge of the contracting side maintains its position despite the continuous addition of new network. This outcome can be explained as follows: once a few anchors are removed locally, the force load on the anchors remaining in the vicinity increases, making their removal in the following cycles more likely than in other areas. At the same time, anchor removal leads to the collapse of the network, concentrating dipoles and further increasing the force load. These mechanical events effectively generate a mutual negative feedback between adhesion and contraction – a mechanism postulated in several previous studies [33, 48]. Our simulations demonstrated that stochasticity of local dynamics eventually leads to self-organized growing and stable edges segregated to opposite sides of the system, producing polarized distributions of dipoles and anchors. Dipoles become concentrated towards the interior of the system, with the region of highest density shifted to the contracting side, while the anchors are present along the entire perimeter, but are internalized into the bulk only at the growing side where they are not removed (Figures 4J and 4L).

#### Migrating

Finally, a migrating phenotype is observed at intermediate ARTF ratios between 3 and 10, depending on the anchor density (lower ARTF ratios require higher anchor density). In contrast to growing phenotypes, the size of these systems reaches a steady state, and they move persistently several diameters before eventually changing direction (Figure 5B, Supplementary Movie S5). The dynamics of these systems is similar to anisotropic growth: anchors are removed at one side (trailing edge) and persist at the other side (leading edge), the network displays strong inward flow at the trailing edge and low and disordered flow at the leading edge, and the distribution of dipoles and anchors is strongly polarized. Unlike anisotropically growing systems, contraction at the trailing edge not only balances the addition of new network but leads to a steady translocation of the whole system. Despite uniform addition of network and anchors everywhere at the periphery, anchors added at the back are quickly removed and the system robustly maintains its polarity (Figure 5F, inset). The shape and dynamics of these systems bear a strong resemblance to persistently migrating cells like fish and amphibian keratocytes: the system is elongated perpendicularly to the direction of motion, the leading edge is convex, and the trailing edge, concave, the network and dipoles are concentrated in a bundle-like structure running parallel to the trailing edge, anchors are found at the front and sides, and dipoles at the back. Similar to a keratocyte, the system advances steadily at the front while displaying variable edge velocity at the back characterized by short-lived protrusions followed by rapid retraction [1, 49] (Figure 5F, inset.

**Fig 5.**
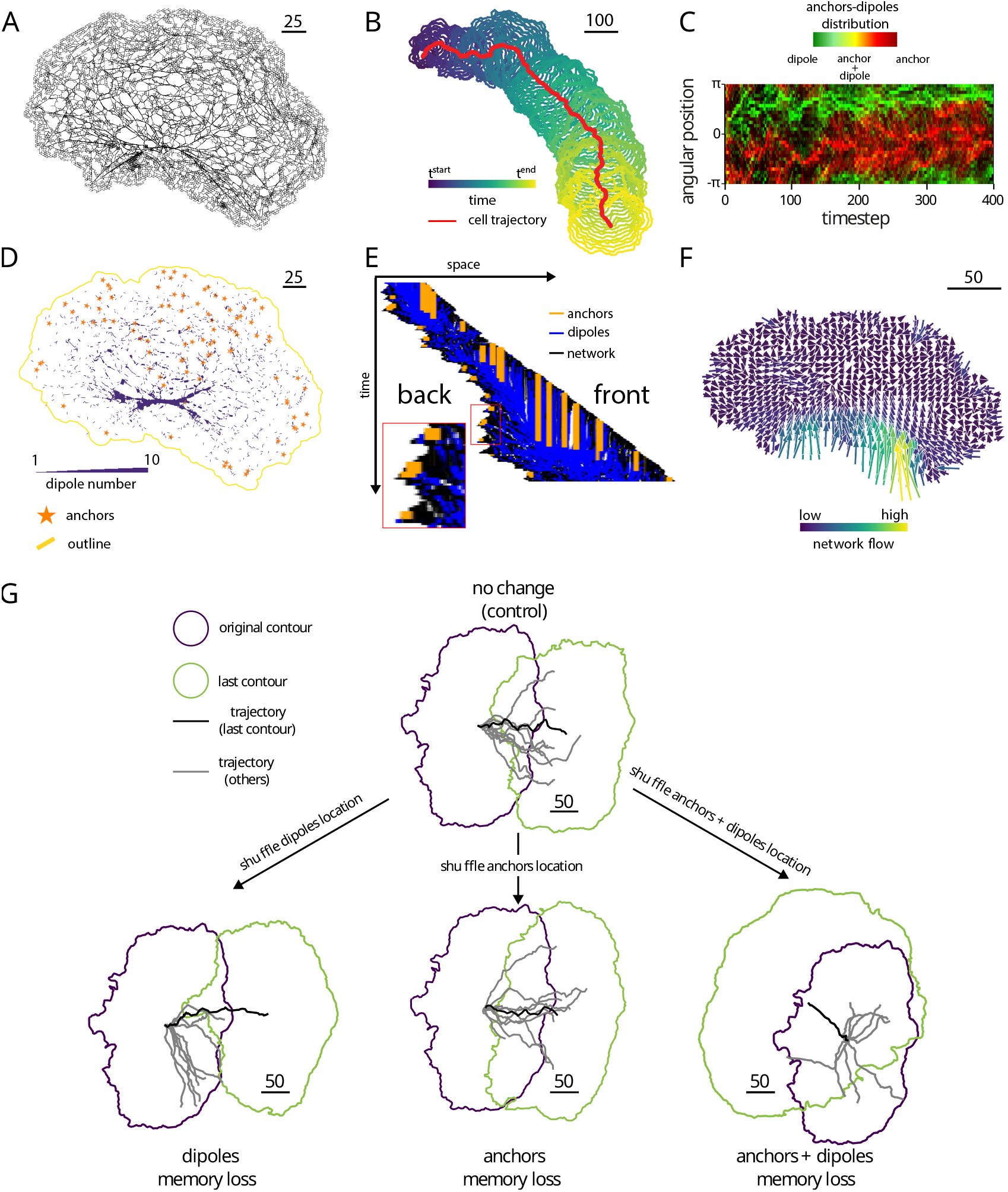
Intermediate ARTF ratio leads to spontaneous polarization and persistent migration. Features and characteristic movement of a system with intermediate ARTF ratio. A) Network configuration. B) Movement of the system throughout the simulation (400 steps). The position of the centroid is drawn in red. The system outline is shown every 10 steps, color coded for time from dark blue to yellow (viridis palette). C) Circular kymograph of dipoles (green) and anchors (red) (see methods). D) Distribution of anchors (orange stars) and dipoles (blue) with the system outline (yellow). The width of dipoles represents the number of overlapping dipoles. E) Kymograph along a line going through the center of the system. Network is represented in black, dipoles in blue, and anchors in orange. Inset: Zoom on the back of the cell showing multiple short-lived protrusion-retraction events F) Network displacement before and after minimization. Amplitude of the displacement is color coded from dark blue (low flow) to yellow (high flow). G) Effect of randomization of anchor and/or dipoles on a polarized system. Top panel shows 20 simulations of the control case (without any randomization) with a representative initial (resp. last) configuration in blue (resp. green) and the corresponding cell trajectory in black. Other trajectories are displayed in gray. The three lower panels represent respectively spatial perturbation of dipoles (left), anchors (middle) and both dipoles and anchors (right). The scale bar is given in model distance units.

To probe the mechanism of polarization in the migrating phenotype, we tested if the direction of motion is determined by the polarized distribution of anchors, dipoles, or both. To this end, at one of the steps of the simulation where the system displayed directional motion, we randomized the distribution of either anchors, or dipoles, or both and ran multiple realizations of the simulation for each case. Resulting trajectories were compared to multiple realizations of the same simulation without randomization of the elements. We observe that after randomization of just one type of the elements, the system displays a directional bias indistinguishable from the system without randomization, and only the randomization of both anchors and dipoles results in a loss of directionality (Figure 5). This result is consistent with the idea of a feedback between adhesion and contraction as described above.

### Intermediate anchor lifetime is optimal for motion

The above observations suggest that anchor turnover is important for the type of system behaviour. To characterize anchor turnover in quantitative terms, we measure mean anchor lifetime and investigate how it depends on the ARTF ratio and anchor density (Figure 6A). Anchor lifetime increases rapidly with the ARTF ratio, and the increase happens earlier with higher anchor density. These results can be explained as follows. Over multiple simulation steps, bonds and dipoles tend to form bundled structures in which several dipoles are aligned. These structures, reminiscent of stress fibers, are generally found between anchors. The stacking of actively pulling network elements in parallel leads to an increase in the force on the anchor that eventually reaches its threshold and the anchor is removed. The ARTF ratio sets the minimal number of dipoles that must pull on a single anchor to remove it. As a result, the anchor lifetime is correlated with this ratio; the greater the ratio, the more dipoles are required to remove an anchor and therefore the more steps are required to build a structure with sufficient pulling force. The anchor density has an impact on the anchor lifetime as well: a larger density implies that the force is distributed among a larger number of anchors, and it takes longer to reach the threshold. Eventually, at high a ratio of anchor removal threshold to dipole force, large structures are not able to remove anchors fast enough to balance the addition of new ones, and the systems grow uncontrollably. Note that the measurement of anchor lifetime appears to reach a finite plateau. This is an artifact due to the limit on the size of the system. In systems with a ratio of 1000, no anchor is removed from the system and therefore anchors have virtually infinite lifetimes.

**Fig 6.**
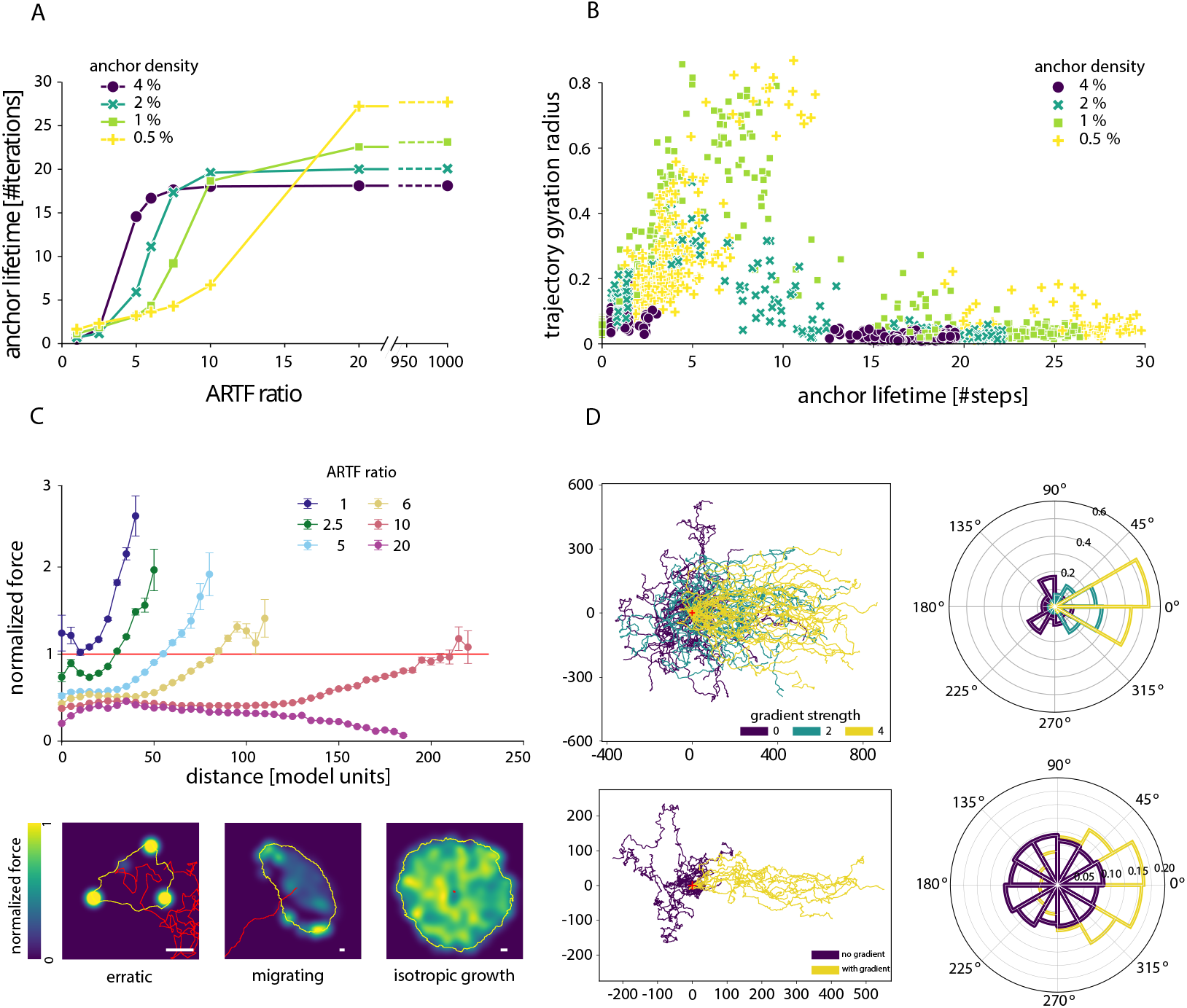
Intermediate anchor lifetime is optimal for migration. A) Plot of anchor lifetime as a function of the ARTF ratio for different values of the anchor density. Every point is an average over 10 simulations. B) Scatter plot of trajectory gyration radius as a function of the anchor lifetime for different values of the anchor density. Each point is a single simulation. C) (top) Force-distance relationship for different values of the ARTF ratio averaged over 10 simulations. Force is normalized by the anchor removal threshold. Distances are discretized in bins of 5 model length units from the system centroid (defined as the centroid of the system outline). Anchor density is 1%. Bottom panel, spatial force distribution for typical realization of erratic (left), isotropic growth (center) and migrating (right) phenotypes. The corresponding cell trajectories and outlines are shown in red and yellow. Amplitude of the normalized force is color coded from dark blue (low forces) to yellow (high forces). D) Effects of ARTF ratio gradient strengths for cells with migrating (top) and erratic (bottom) cell phenotypes. Trajectories are displayed for different gradient values, see methods, (left panels) together with their trajectories’ end-to-end vectors angular distributions (right panels).

Anchor lifetime, in turn, is a determining factor for the motion of the system. We characterized the extent of motion of the system by the gyration radius of the trajectory (see methods). Figure 6B shows that the gyration radius peaks at intermediate anchor lifetime of around 5 simulation steps. Here, the systems are observed that move in a relatively persistent fashion. Shorter lifetimes mean that the anchors are removed too frequently for the system to develop any memory and polarize. This results in a smaller gyration radius and corresponds to the erratically moving systems and systems with high centripetal flow. Above a lifetime of 5, the gyration radius decreases again. Anchors live too long and become a hindrance to motion until the systems do not move at all (anisotropically and isotropically growing systems).

### In polarized systems, forces increase with the distance from the center and exceed anchor removal threshold only at the periphery

In our recent work [2], we showed that local stress patterns exerted by rapidly moving fish epidermal keratocytes correlate with distance from the cell center and that edge retraction initiates preferentially at the longest distances, leading eventually to emergence and maintenance of a characteristic cell shape with long axis perpendicular to the direction of motion [1]. Using a simple mechanical model of the actin-myosin network without turnover we showed that the increase of force with distance from the center is intrinsic to such networks. Here we test whether a system with turnover exhibits a similar feature and how it depends on anchor turnover parameters (Figure 6C, Supplementary Movies S6-S8). Systems that build forces above the anchor removal threshold display increased force with distance from the center. When the system cannot build enough force to remove the anchors, this pattern is lost, and force instead decreases with distance as in an isotropically growing phenotype (Figure 6C, ratio 20, Supplementary Movie S7). Force-distance curves for polarized systems show forces large enough to remove anchors only at the system periphery (Figure 6C, ratio 5 to 10, migrating and anisotropically growing systems, Supplementary Movie S8), consistent with our findings in fish keratocytes [2]. In contrast, erratic and centripetal systems (Figure 6C, ratio 1 and 2.5, Supplementary Movie S6) generate forces above anchor removal threshold everywhere (Figure 6C, ARTF ratio 1). Anchor turnover at each step prevents these systems from building polarity.

### External cues can bias migration

As shown above, the system of bonds, dipoles, and anchors is able to polarize and move spontaneously. We next tested if an external directional cue could bias this system. To simulate an external cue, we introduced a directional gradient in the ARTF ratio. Such a gradient could reflect the input of a substrate signal, e.g. gradient of ligand density or rigidity (haptotaxis or durotaxis).Addition of an ARTF ratio gradient clearly biases the direction of migration of polarized systems towards higher ARTF ratio in a manner dependent on the strength of the gradient (Figure 6D and Supplementary Movie S9), and also induces directional motion in the centripetal system that does not migrate in the absence of a gradient (Figure 6D and Supplementary Movie S10).

## Discussion

Here, we found that a minimalistic model system consisting only of elastic bonds, force dipoles, and anchors, subjected to very simple turnover rules, is capable of self-organized polarization and directional motion. The key feature of this model is the detachment of anchors upon the buildup of a threshold force. The timescale of this detachment effectively defines the behaviour of the system. If anchors detach at every simulation step, the system cannot build up any form of memory, fails to self-organize, and moves erratically, while if anchors are effectively permanent, the system expands isotropically. Critically, only when anchors detach at an intermediate rate can the system build up asymmetry, spontaneously polarize and undergo directed motion.

This result suggests that the ability to self-polarize is an intrinsic property of such an active mechanical network. Importantly, symmetry breaking results from mutual mechanics of contraction and adhesion. Our model features the addition of new network, mimicking actin protrusion, but the addition is uniform around the perimeter of the system and thus does not contribute to symmetry breaking. The system polarizes despite uniform protrusion. Likewise, our model does not feature any feedback from the outer boundary, which could represent membrane tension, and therefore such feedback is not necessary for symmetry breaking. Neither does the model feature any mass conservation constraints, thus excluding any mechanisms that depend on a limited supply of components. While all the above mechanisms could be a part of the polarization process in living cells, our study suggests that the self-organization of contraction and adhesion alone is sufficient to break symmetry. This could be a key triggering event, especially in cases where polarization starts by local retraction at the prospective rear [19, 20]. Asymmetry of protrusion may develop subsequently due to a limited actin pool or other mechanisms. Additional features may also result from adhesion dynamics that in reality are richer than represented in our model, such as reinforcement under applied force and viscous stick-slip behavior [34, 50, 51].

The switch from isotropic behavior to polarization happens in our system through tuning the balance between contractility and adhesion. This is consistent with many experimental studies indicating that intermediate adhesion strength and an an optimal balance between contractility and adhesion favor fast cell migration [52, 53]. However, our results show that optimal balance between contractility and adhesion does not just speed up cell motion, but itself promotes the development of polarity. This suggests that any factor influencing contractility or adhesion may also promote or suppress symmetry breaking. This finding provides a framework for understanding the roles of various regulatory factors and molecules in cell polarization and suggests a possible intersection between chemical signaling networks and cytoskeletal machinery. According to our model, a trigger for polarization does not have to be a chemical gradient or other directional signal. Instead, a global change in the chemistry of contractile or adhesive machinery may trigger symmetry breaking. A similar finding was recently reported for a different system - a biomimetic model based on actin assembly [54]. At the same time, our simulation of the imposed spatial gradient of the adhesion strength demonstrated that the machinery is sensitive to directional signals, which can both promote polarization and influence the direction of motion in a system that is already polarized. An additional implication of these findings is that the signal-induced symmetry breaking and the cue providing directional information could be separate and independent signals.

The polarization mechanism in our model is consistent with the geometrical idea of distance-dependent protrusion-retraction switch and with the experimentally observed distance-dependence of forces [1, 2]. Forces in the system build up at the most distant anchor points, eventually leading to their detachment. This limits lateral expansion of the system and results in a geometry reminiscent of fish keratocytes with a long axis normal to the direction of motion. This polarization process is also consistent with the idea of feedback between motion and polarity, such as in UCSP model [30] : as the system moves, force dipoles accumulate at the back leading to the force build-up and detachment of adhesions, which promotes continued motion in the same directions, while adhesions at the front remain through multiple simulation cycles. Furthermore, the spatial and temporal scales of the model are quantitatively consistent with those of real cells. Because force dipoles act between neighbouring network nodes, the distance between nodes (2 model units) can be mapped to the characteristic length of a myosin bipolar minifilament, approximately 300nm [37]. Considering that the size of moving systems in our simulations is in the range of tens of thousands square model units, we arrive at system linear dimensions in the range of tens of micrometres, comparable to those of migrating cells. The protrusion rate in the model is fixed at 4 model units per simulation step (~600nm). Given typical cell protrusion rates of several tens to a few hundred nanometres per second [55, 56], this implies a simulation time step on the order of seconds. Accordingly, the full simulation duration (400 steps) corresponds to tens of minutes (typically 40min for a protrusion rate of 100nm/s), consistent with the timescale required for cell polarization and migration over distances of several cell lengths. Thus, our model provides realistic relations between the microscopic processes and system-level behaviour in both space and time. Importantly, realistic system dimensions are not set but emerge spontaneously in the model through self-organization.

## Conclusion

By dissecting polarization mechanisms via physical modeling, our study thus identifies network of elastic, contractile, and adhesive elements as an independent module capable of symmetry breaking. Experimental biomimetic studies reconstituting active systems from purified components provide another important approach to dissecting polarization mechanisms. These studies mainly focused on reconstituting actin networks at the surface of small movable objects or inside lipid vesicles [54, 57–60]. Our work demonstrates that feedback from the external boundary is not a necessary part of the polarization mechanism and suggests that symmetry breaking could be achieved in patches of actin-myosin network by tuning their attachment to a flat surface. This could inspire new biomimetic studies.

In living cells, symmetry breaking through contraction-adhesion mechanics is probably but one of multiple polarization mechanisms. Even this mechanism is likely much more complicated than represented in our model due to phenomena such as adhesion reinforcement, viscous dissipation of forces, etc. Nevertheless, by identifying a simple relationship between adhesion lifetime and polarization, our study provides a framework for understanding the integration of different mechanisms in cell polarization and a benchmark for further experimental and theoretical studies.

## Supporting information

### Additional resources

1. zenodo repository: DOI:10.5281/zenodo.16925923
2. gitlab repository: https://gitlab.unige.ch/Zeno.Messi/balanced-contractility-and-adhesion-drive-polarization-in-a-minimal-elastic-actomyosin-network

### Supplementary Figures

### Supplementary Videos

**Movie S1. Erratic system timelapse**. Timelapse of a typical simulation of the erratic phenotype. Bonds are shown as black segments black, dipoles as blue segments, and anchors as orange stars. Trajectory of the system centroid from the start of the simulation to the current position is shown in red. Grid size 50 *×* 50. Inset shows the outline of the system in black and the trace of the centroid in red. Reference for the simulation 1711312861 (same as Figure 3A-F). Length of the simulation: 400 steps.

**Movie S2. Centripetal system timelapse**. Timelapse of a typical simulation of the centripetal phenotype. Bonds are shown as black segments black, dipoles as blue segments, and anchors as orange stars. Trajectory of the system centroid from the start of the simulation to the current position is shown in red. Grid size 50 *×* 50. Inset shows the outline of the system in black and the trace of the centroid in red. Reference for the simulation 1708960333 (same as Figure 3G-L). Length of the simulation: 400 steps.

**Movie S3. Isotropic growing system Timelapse**. Timelapse of a typical simulation of the isotropic growing phenotype. Bonds are shown as black segments black, dipoles as blue segments, and anchors as orange stars. Trajectory of the system centroid from the start of the simulation to the current position is shown in red. Grid size 50 *×* 50. Inset shows the outline of the system in black and the trace of the centroid in red. Reference for the simulation 1707691522 (same as Figure 4A-F). Length of the simulation: 400 steps.

**Movie S4. Anisotropic growing system Timelapse**. Timelapse of a typical simulation of the anisotropic growing phenotype. Bonds are shown as black segments black, dipoles as blue segments, and anchors as orange stars. Trajectory of the system centroid from the start of the simulation to the current position is shown in red. Grid size 50 *×* 50. Inset shows the outline of the system in black and the trace of the centroid in red. Reference for the simulation 1711535798 (same as Figure 4G-L). Length of the simulation: 400 steps.

**Movie S5. Migrating system timelapse**. Timelapse of a typical simulation of the migrating phenotype. Bonds are shown as black segments black, dipoles as blue segments, and anchors as orange stars. Trajectory of the system centroid from the start of the simulation to the current position is shown in red. Grid size 50 *×* 50. Inset shows the outline of the system in black and the trace of the centroid in red. Reference for the simulation 1710023558 (same as Figure 4G-L). Length of the simulation: 400 steps.

**Movie S6. Force map timelapse of erratic system**. Timelapse of force maps generated by a typical simulation of the erratic phenotype. System outline is shown in yellow. Trajectory of the system centroid from the start of the simulation to the current position is shown in red. Magnitude of the force on nearby anchors is shown color-coded. The force is normalized at each frame by the maximum for on an anchor in the frame. Frame number is shown in the left-top corner. Reference for the simulation 1711312861 (same as Figure 6C (erratic)). Length of the simulation: 400 steps. See methods for a description of the generation of the force maps.

**Movie S7. Force map timelapse of migrating system**. Timelapse of force maps generated by a typical simulation of the migrating phenotype. System outline is shown in yellow. Trajectory of the system centroid from the start of the simulation to the current position is shown in red. Magnitude of the force on nearby anchors is shown color-coded. The force is normalized at each frame by the maximum for on an anchor in the frame. Frame number is shown in the left-top corner. Reference for the simulation 1723538402 (same as Figure 6C (migrating)). Length of the simulation: 400 steps. See methods for a description of the generation of the force maps.

**Movie S8. Force map timelapse of isotropic growing system**. Timelapse of force maps generated by a typical simulation of the isotropic growing phenotype. System outline is shown in yellow. Trajectory of the system centroid from the start of the simulation to the current position is shown in red. Magnitude of the force on nearby anchors is shown color-coded. The force is normalized at each frame by the maximum for on an anchor in the frame. Frame number is shown in the left-top corner. Reference for the simulation 1707691522 (same as Figure 6C (isotropic growth)). Length of the simulation: 400 steps. See methods for a description of the generation of the force maps.

**Movie S9. Trajectory comparison with and without directional cue for migrating systems**. Comparison of trajectories of systems with different level of external cue. (Top) no cue, (middle) gradient 2, (bottom) gradient 4. Outline and trajectory of each system is shown in black and red respectively. All simulations were generated from the same set of parameters that corresponds to the migrating phenotype.

**Movie S10. Timelapse of a centripetal system with directional cue**. Timelapse of a system from the centripetal phenotype with an external cue. Bonds are shown as black segments black, dipoles as blue segments, and anchors as orange stars. Trajectory of the system centroid from the start of the simulation to the current position is shown in red. Grid size 50 *×* 50. Inset shows the outline of the system in black and the trace of the centroid in red. Reference for the simulation 1743622318. Length of the simulation: 400 steps.

## Acknowledgments

The work was supported by the National Science Foundation (A.B.V.), Switzerland via the grant 31003A 169972, the European Molecular Biology Organization via the EMBO Postdoctoral fellowship grant EMBO ALTF 1053-2022 (Z.M.), and the Francis Crick Institute (N.W.G.), which receives its core funding from Cancer Research UK (CC2119), the UK Medical Research Council (CC2119), and the Wellcome Trust (CC2119).

## Author contributions

Conceptualization, Z.M. and F.R. and A.B.V.; methodology, Z.M. and F.R. and A.B.V.; investigation, Z.M. and F.R. and A.B.V.; data analysis, Z.M.; writing-–original draft, Z.M. and A.B.V.; writing-–review & editing, Z.M. and F.R. and A.B.V. and N.W.G; funding acquisition, Z.M. and F.R. and A.B.V. and N.W.G.; resources, F.R. and N.W.G. and A.B.V.; supervision, F.R. and A.B.V.

## Notes

https://zenodo.org/records/16925923

https://gitlab.unige.ch/Zeno.Messi/balanced-contractility-and-adhesion-drive-polarization-in-a-minimal-elastic-actomyosin-network

### Competing Interest Statement

The authors have declared no competing interest.

### Summary of Updates

Manuscript revised; Material and Methods revised; New figure 2.

